# Squid Ring Teeth Coated Mesh Improves Abdominal Wall Repair

**DOI:** 10.1101/214114

**Authors:** Ashley N. Leberfinger, Monika Hospodiuk, Abdon Pena-Francesch, Bugra Ayan, Veli Ozbolat, Srinivas Koduru, Ibrahim T. Ozbolat, Melik C. Demirel, Dino J. Ravnic, DO

**Author notes:** Corresponding Authors: Dino J. Ravnic, DO, MPH, Penn State Health Milton S. Hershey Medical Center, Department of Surgery, Division of Plastic Surgery, 500 University Drive, Hershey, PA 17033; Melik C. Demirel, PhD, Center for Research on Advanced Fiber Technologies (CRAFT), Materials Research Institute & Huck Institutes of Life Sciences, Department of Engineering Science and Mechanics, Pennsylvania State University, University Park, PA, 16802.

## Abstract

**Background:** Hernia repair is a common surgical procedure with mesh often used. Current mesh materials have a high incidence of repair failures, due to poor tissue integration, and complications such as seroma and pain. Polypropylene (PP) mesh is the standard material in hernia repair secondary to its material durability; however, failures still approach 15%. In this first time animal study, we hypothesized that squid ring teeth (SRT), a biologically-derived high strength protein, coated polypropylene (SRT-PP) mesh, would offer enhanced tissue integration and strength compared to standard PP mesh, while proving biocompatibility for *in vivo* use.

**Materials and methods:** Polypropylene mesh was coated with SRT. Mechanical properties and cell proliferation studies of the composite mesh were performed *in vitro.* Rats underwent inlay mesh implantation in an anterior abdominal wall defect model. Repair was assessed clinically and radiographically, with integration evaluated by histology and mechanical testing.

**Results:** Cell proliferation was enhanced on SRT-PP composite mesh. This was corroborated by abdominal wall histology, dramatically diminished cranio-caudal mesh contraction, improved strength testing, and higher tissue failure strain following *in vivo* implantation. There was no increase in complications with SRT, with regard to seroma or visceral adhesion. No foreign body reactions were noted on liver histology.

**Conclusion:** SRT-PP mesh showed better tissue integration than PP mesh. SRT is a high strength protein that is applied as a coating to augment mesh-tissue integration leading to improvements in abdominal wall stability with potential to reduce re-intervention for failures.

## INTRODUCTION

Hernia repair is one of the most common surgical procedures worldwide (1). Approximately 400,000 procedures are performed annually in the United States for correction with associated costs exceeding three billion dollars per year (2). A hernia can occur at any point of weakness of the abdominal wall, but are especially likely to develop at the site of a previous fascial incision (3). Development of a hernia can be associated with pain, discomfort, and weakness leading to significant lifestyle alteration (4). Furthermore, hernias can result in intestinal obstruction and strangulation requiring emergency surgery leading to associated morbidity and mortality (5).

Reparative surgery utilizing mesh for abdominal wall repair or reinforcement is often indicated, especially in complicated hernia repairs (6). It is estimated that one-third of patients with a recurrent ventral hernia repair experience an additional failure, and with each subsequent repair, there is a step-wise worsening of long-term outcomes (7). Since Usher widely introduced a plastic prosthesis for hernia repair in 1955, non-absorbable synthetic mesh, such as polypropylene (PP) has become widespread (8). In fact, it has become a mainstay of hernia repair worldwide as it decreases hernia recurrences compared to primary closure (9). However, its intraperitoneal use has been questioned since Kaufman et al. first documented fistula formation in 1981(10) and has led to legal concerns more recently (11). This has led to the development of biologic and composite mesh, especially for use in patients which an overlay or sublay repair cannot be performed and the implanted material is in direct contact with the viscera. Biologic mesh is derived from mammalian tissue such as human dermis (Alloderm), porcine dermis (Strattice), or bovine pericardium (Veritas). When placed in an inlay or bridge position, it offers visceral protection while minimizing the risk of fistula formation, infection, and other wound complications, especially in a contaminated field (12). Despite these benefits, the average cost of an initial abdominal wall reconstruction with a biological mesh exceeds $85,000 per patient (13). Furthermore, the use of biologic mesh in a bridged position leads to recurrence rates >60% and reoperation in >70% of patients (14). This has led many practitioners to utilize an absorbable mesh with a planned second procedure in patients where initial fascial closure is not possible (14). Alternatively, some surgeons have turned to composite mesh when faced with this scenario. Composite meshes typically consist of a PP or polyester base, which are covered with a protective membrane/film to prevent against visceral adhesion or fistula formation (15). However, by limiting tissue integration these coatings may limit the strength of the final repair and expectedly are more reliable when used in an underlay position (16). While advanced surgical techniques, such as anterior (17) or posterior (18) component separation, have improved outcomes, hernia recurrence can still approach 15% in the most complicated cases, despite the use of mesh in any position (19). Furthermore, when fascial closure is not possible, the unavailability of a mesh that performs well in the inlay or bridging position has led towards the investigation of total abdominal wall transplantation (20). Therefore, a major materials challenge is to develop an affordable material, which can be utilized to provide a strong and definitive repair in patients where fascial re-approximation is not possible.

Our unique solution to this problem is a structural protein, which offers high strength and biocompatibility. Squid ring teeth (SRT), found both in the arms and tentacles of squid have high elastic modulus (2-4 GPa) and excellent mechanical strength (50-100 MPa) (21). The rings are composed of a group of repetitive proteins, ranging from 15 to 60 kDa, which have a segmented sequence topology similar to semi-crystalline segmented copolymers (22, 23). The repetitive sequence favors the formation of a beta-sheet-rich cross-links that confers unique mechanical (24), adhesive (25), thermal (26), optical (27), and self-healing (28) properties to SRT. Furthermore, SRT is easily processed by solution and thermal fabrication methods into nano to macro protein-based materials, from nanotubes and nanoparticles to large-scale objects (25). However, SRT has never been used in animal studies earlier. The objective of this study is to evaluate the applicability of SRT protein for soft-tissue repair in an abdominal wall defect model. We hypothesized that SRT would offer superior abdominal wall integration and strength when used as a coating compared to standard PP mesh, while proving to be safe for animal implantation.

## MATERIALS AND METHODS

### Fabrication and Characterization of Squid Ring Teeth Protein-coated Hernia Mesh

Squid ring teeth were extracted from suction cups in the arms and tentacles of *Loligo Pealeii* squid (Figure 1a-c). The rings are composed of a group of proteins ranging from 15 to 60 kDa (Figure 1d). SRT samples were washed in water and ethanol, and then dried in ambient conditions overnight. They were dissolved overnight in 1,1,1,3,3,3-Hexafluoro-2-propanol (HFIP) to a concentration of 50 mg/mL and the solutions were purified by centrifugation. Polypropylene mesh (Bard; New Providence, NJ) was cut into 2×5 cm strips and each strip was dip coated with SRT/HFIP solution (Figure 1e). SRT/HFIP has been extensively used with polymers (29, 30) and proteins (31), and has been recently used with SRT proteins in the fabrication of composites (32). The coated strips were dried at room temperature (RT) and washed in deionized water to remove any residual HFIP solvent. The resulting SRT-coated PP mesh strips (SRT-PP) had 15±5 % of protein content (w/w). Spectral data was collected (Thermo Nicolet IR) under attenuated total reflection ATR (diamond crystal) mode using Norton-Beer apodization with 4 cm-1 resolutions. For each spectrum, 256 scans were co-added. Uniaxial tensile testing of composite SRT-PP and uncoated PP mesh (control) was performed in a TA 800Q DMA instrument. Thin film fixtures were used to clamp the specimens and strain ramp measurements were performed at a strain rate of 5 %/min.

**Figure 1.**
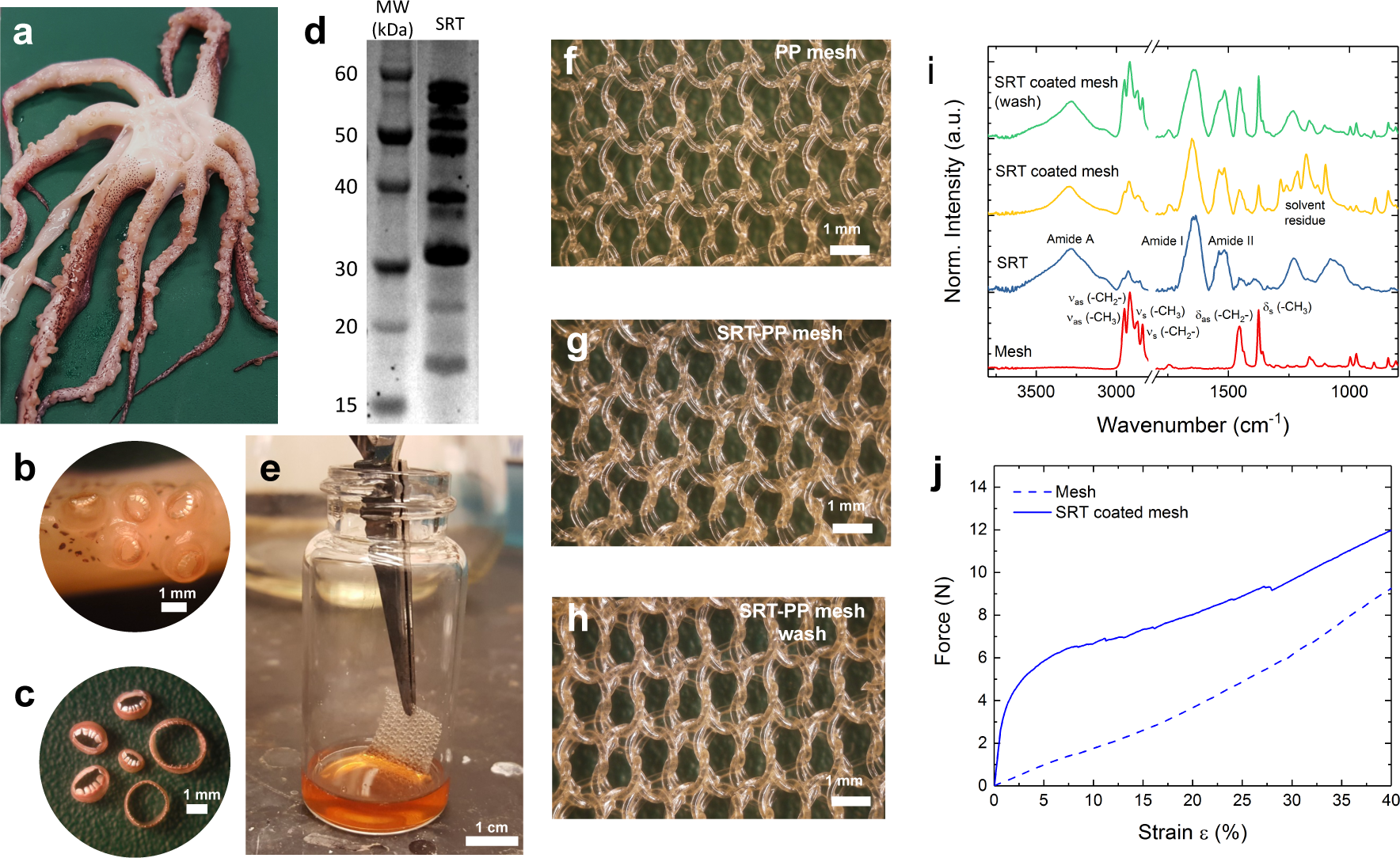
SRT protein and *in vitro* testing of SRT mesh. (a) Common squid *Loligo Pealeii*, (b) suction cups on tentacles, and (c) SRT. (d) SDS-Protein gel for SRT. (e) Dip coating of PP mesh in SRT/HFIP solution. (f) Uncoated PP mesh. (g) Composite SRT-PP mesh after coating. (h) Composite SRT-PP mesh after washing (removal of solvent). (i) FTIR spectrum of PP mesh, SRT protein, composite SRT-PP mesh, and composite SRT-PP mesh after washing. (j) Uniaxial tensile testing of composite SRT-PP and control PP mesh.

### Cell Preparation and Adhesion Assay

NIH/3T3 fibroblasts (ATCC; Manassas, VA) were cultured in Dulbecco’s Modified Eagle’s Medium (DMEM, Corning Cellgro; Manassas, VA) supplemented with 10% FBS (Life Technologies; Grand Island, NY), 100U/mL Penicillin G, and 100 µg/mL streptomycin (Life Technologies). Cells were maintained at 37°C in 5% CO_2_. Sub-confluent cultures were split using 0.25% trypsin-0.1% EDTA (Life Technologies). Meshes were soaked in 70% ethanol for five minutes and washed three times in sterile phosphate buffered saline (PBS), then once in sterile 3T3 media, and placed into individual wells of an untreated 24 well plate. 3T3 cells were seeded at a concentration of 2×10^6^ cells suspended in 300 µL 3T3 media. The plate was incubated for six hours at 37°C, then an additional 150 µL of media was added and incubated overnight. The mesh was then transferred to new wells in a cell-repellent plate. Cell culture medium was changed every two days over a seven-day period. Meshes were imaged with an EVOS FL Auto inverted microscope (ThermoFisher, Pittsburgh, PA). Meshes were stained to determine adhesion at three time points (one, four, and seven days). At each time point, three SRT-PP and three PP meshes were rinsed three times with Dulbecco’s Phosphate Buffered Saline (DPBS; Life Technologies). 400 µL DPBS with 2 µM calcein-AM (Invitrogen; Carlsbad, CA) and 4 µM ethidium homodimer (Life Technologies) was then added to the wells. Plates were protected from light and incubated at 37°C in 5% CO_2_ for 30 minutes, then rinsed three times with DPBS. Four representative areas of each SRT-PP and PP mesh were randomly selected for imaging at each time point. Adhesion was quantified using ImageJ (National Institutes of Health; Bethesda, MD). Briefly, the green channel was converted to a binary image, evaluated using threshold adjustment, and counted in ImageJ. The live cell number was divided by the area of the mesh to obtain cells/mm^2^. After seven days of culture, meshes were fixed overnight with 4% paraformaldehyde (Sigma-Aldrich; St. Louis, MO) at 4°C. Constructs were washed three times in DPBS at RT. Permeabilization was performed with 0.25% Triton X-100 (Sigma-Aldrich) and 1% bovine serum albumin (RPI Corp.; Mount Prospect, IL) diluted in DPBS and incubated for one hour. Meshes were double stained with Alexa Fluor 568 phalloidin diluted 1:100 in DPBS and 5 µg/mL DAPI for one hour. Samples were washed three times per 10 min in DPBS and imaged on a confocal laser scanning microscope (Olympus FV10i; Tokyo, Japan).

### Scanning Electron Microscopy (SEM)

Field emission scanning electron microscopy (SEM; Nova NanoSEM 630, FEI; Hillsboro, OR) was used to investigate surface topography. After seven days of culture, meshes were fixed overnight with 4% paraformaldehyde (Sigma-Aldrich) at 4°C. Constructs were washed in DPBS at RT and dehydrated using graded ethanol solutions (25% to 100%). Meshes were further dried in a critical point dryer (CPD300, Leica EM; Wetzlar, Germany). After complete dehydration, meshes were sputter coated with iridium (K557X Emitech Sputter Coater, TX), 8 nm thickness, and observed at an accelerating voltage of 10 keV through a detector.

### Vertebrate Animal Procedures

All animals are monitored twenty-four hours a day by appropriately trained personnel from the Department of Comparative Medicine under the supervision of the Dr. Ravnic, as well as the project veterinarian. The animals are housed individually in specially designed cages, which allow freedom of movement but preclude injury or discomfort to the animal. Animal studies are performed in AAALAC-accredited animal facilities operated by the Department of Comparative Medicine. Veterinary care is provided under the direction of the veterinary staff of the Department of Comparative Medicine, and is administered in accordance with guidelines set forth in The Guide for Care and Use of Laboratory Animals, 8th edition. In Penn State facilities staff veterinarians are available for consultation twenty-four hours a day, seven days a week. All necessary health care, nutritional necessities, and husbandry requirements for these animals is provided through the Department of Comparative Medicine in accordance with recommendations set forth in the Guide, as well as all applicable state, federal, and local regulations.

### In Vivo Implantation of SRT-coated Mesh

Surgical anesthesia and postoperative analgesia is administered by Dr. Ravnic under the supervision of the project veterinarian. Under Penn State institutional animal protocol #47197, nine 10-week-old Sprague Dawley rats (four male and five female) were anesthetized with inhalational isoflurane and a subcutaneous injection of carprofen (5mg/kg) was administered. A warm water blanket was used to prevent excessive heat loss and aseptic conditions were used. The rat was then moved to the surgical field and a sterile drape was applied. Ethylene oxide was used to sterilize the SRT mesh. A 2×5 cm full-thickness segment of the anterior abdominal wall was excised to allow for mesh placement. The fascia and muscle layers were repaired with an inlay (bridging) technique. Three males and three females received SRT-PP mesh whereas two females and one male received PP mesh. Mesh was secured to the abdominal wall using 4-0 monocryl sutures in an interrupted fashion. Following mesh placement, the overlying skin was closed with 4-0 monocryl sutures in a subcuticular fashion. Postoperatively animals were examined daily for the first week then three times per week thereafter for weight loss, pain, hernia development, or other complications.

### In Vivo Magnetic Resonance Imaging

At day 60, animals underwent Magnetic Resonance Imaging (MRI). Inhalational isoflurane gas was used for anesthesia. Animals were placed inside an animal holder (Acrylic Glass^®^ animal cradle equipped with a nosecone and adjustable bite bar) and then into the MRI (7-Tesla 300MHz 70/20as Bruker Biospec MRI system, Bruker Biospin; Ettlingen, Germany). A respiration monitor (PC-SAM Model 1025, SA Instruments; Stony Brook, NY) was used and isoflurane was adjusted in response to the animals’ monitored breaths per minute. After the initial localizer scan was performed, a 2D T2-weighted scan (RARE) was acquired with the following parameters: TR/TE=4101/12.0 ms, 8 averages, 1 echo, 256×256 matrix, 6.0×6.0 cm FOV, RARE factor=8, and slice thickness 1 mm (40 slices). Total acquisition time of the scan (respiratory-gated) was approximately 30 min. Immediately after MRI, anesthetized animals were sacrificed. Image analysis was conducted using ImageJ to detect the presence of any occult hernia development or seroma formation.

### Histological Observations of Tissue Repair and Integration

The anterior abdominal wall was opened widely along three sides of the mesh allowing for hinged exposure of the intra-abdominal contents. Digital photographs were taken and adhesion formation was scored macroscopically. Adhesions were graded using a five point scoring system that accounted for extent (percentage of mesh involved), type (flimsy or dense), and tenacity (how difficult to dislodge) by two blinded observers, as previously described by Deeken and Matthews (33) (Supplemental 1). A 3 cm rim of surrounding abdominal wall was excised along with the mesh. Two specimens from each rat were prepared for histological analysis. The tissue was fixed with 10% formalin and later processed in paraffin, sectioned, and stained with either hematoxylin and eosin (H&E) or Masson’s trichrome. Slides were scored for inflammatory cell infiltrates (neutrophils, eosinophils, macrophages, lymphocytes, and giant cells), neovascularization, necrosis, and hemorrhage as previously described (33) (Supplemental 2). A section of liver was also prepared for H&E staining.

### Biomechanical Testing of Explants

Following macroscopic adhesion assessment, mesh specimens were prepared as 1 cm wide by 5 cm long (3 cm abdominal wall and 2 cm mesh) strips and placed in 0.9% sodium chloride. An Instron 5966 tensile testing device (Norwood, MA) was used to perform uniaxial tensile testing. The specimens were fixed on upper and lower gripers using metal sutures and rigidly held by a 1 kN load-cell platform. The lower gripper held the tissue and the upper gripper held the mesh side of the tissue-mesh explant. Initial thickness and width of the mesh specimens were measured by a digital caliper, then test specimens were manually loaded until positive tension was reached. The length of the reduced section was measured before tensile loads were applied at a loading rate of 500 mm/min until failure was observed. Tensile stress, strain, peak stress, and strain at break were recorded.

## RESULTS

In this study we divided the work into two parts: (1) *in vitro* and (2) *in vivo* testing. The *in vitro* studies showed the strong mechanical properties of the composite mesh and non-toxicity to cells. *In vivo* implantation in rats showed the composite mesh is biocompatible and had superior abdominal wall integration compared to uncoated control. We chose rats for this first time animal study because they are the smallest accepted hernia model in abdominal wall repair studies.

### In Vitro Testing of SRT Mesh

SRT protein is extracted from the arms and tentacles of squid, and cleaned (Figure 1a-c). Molecular weight distribution of SRT protein (15-60kDa) is shown in Figure 1d. The protein is dissolved in an organic solvent for coating PP mesh (Figure 1e). The thickness of the mesh fibers was slightly increased (70 µm to 92 µm) due to the protein coating (15% w/w of protein content) (Figure 1f-h). We verified mesh coating by FTIR (Fourier-transform infrared spectroscopy). FTIR spectrum of PP mesh showed characteristic absorption bands of PP, which are stretching and deformation vibrations of methyl and methylene groups (Figure 1i) (34). SRT showed absorption bands characteristic of proteins or polyamides (22). SRT-PP mesh showed absorption bands from both PP and SRT, indicating that the coating process was successful. There are solvent residues trapped in the film, as shown in the bands in the 1000-1300 cm^−1^ region. Solvent bands were not observed after washing.

Mechanical comparisons were also made between the two groups. PP mesh showed a linear response with strain until failure at 50% strain (unweaving at the edges) (Figure 1j). However, composite SRT-PP mesh showed a clear increase in the tensile load for the same deformation (increase of 6 N), followed by a linear region similar to the PP mesh (Figure 1j). We showed the mechanical data in force units (Newton) instead of engineering stress (Pascal) due to difficulty of estimating the cross sectional area of the mesh.

### Cell Seeding, Count, and Adhesion

NIH/3T3 fibroblasts cells on PP meshes accumulated within the woven areas. Some of the cells aggregated and were delicately attached, which caused loss of these cells during media change. Cells on SRT-PP meshes attached to the SRT coating. Cell density on SRT-PP mesh increased significantly over time, more than threefold on day 7 (8.17 cells/mm^2^ on day 1, 13.02 cells/mm^2^ on day 4, and 29.99 cells/mm^2^ on day 7), while the density on control meshes remained relatively unchanged (around 8 cells/mm^2^) (Figure 2a). The difference between cell content is shown on the confocal images, where cells on composite SRT-PP mesh increased fluorescent intensity compared to PP control mesh (Figure 2b-g).

**Figure 2.**
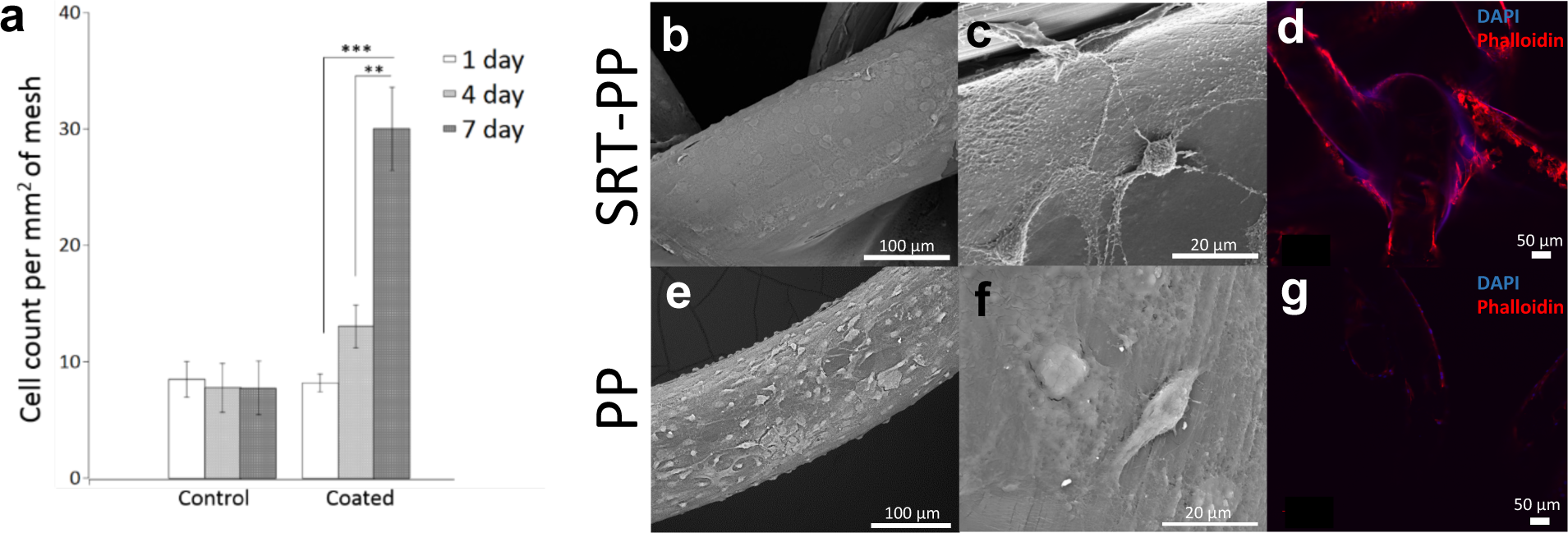
Cell seeding, count, and adhesion. (a) cells/mm^2^ mesh after 1, 4, and 7 days. (b) SEM of SRT-PP mesh. (c) Higher magnification of SRT-PP mesh. (d) Immunofluorescence with DAPI and Phalloidin of SRT-PP mesh. (e) SEM of PP mesh. (f) Higher magnification of PP mesh. (g) Immunofluorescence with DAPI and Phalloidin of PP mesh.

### In Vivo Implantation and Radiographic Evaluation

In our studies all rats had appropriate weight gain (Supplemental 3) and no systemic signs of distress following mesh implantation, indicating biocompatibility of the material. We did not observe any clinical evidence of infection, seroma, or hernia development during the implantation period. MRI of abdominal cavity for composite and control mesh studies showed good mesh incorporation into the abdominal wall and no seromas or occult hernias (Figure 3).

**Figure 3.**
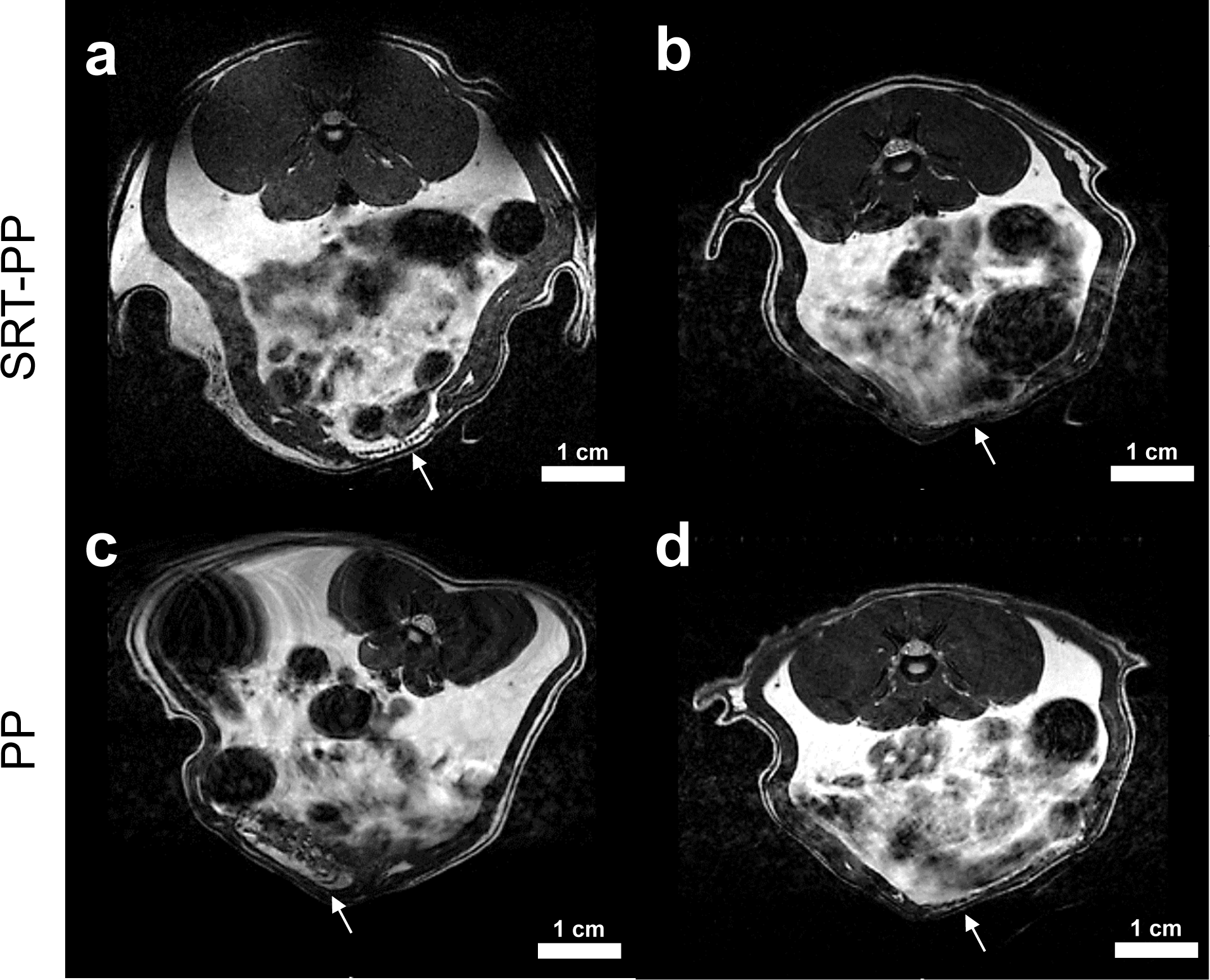
MRI. Good mesh incorporation into the abdominal wall and no seromas or occult hernias in both the SRT-PP (a&b) and PP (c&d) groups. Arrows are pointing to mesh.

### Necropsy and Adhesion Scoring

We observed that upon necropsy, the mesh repair was completely intact in both the SRT-PP and PP control group. However, the control group (N=3) had significant cranio-caudal mesh contraction whereas the SRT-PP group (N=6) had no contraction (2.27 cm vs 0.08 cm, p=0.05) (Figure 4a&e). Despite the three-fold increase in mesh contraction, there was no statistically significant difference in visceral adhesion formation between SRT-PP and PP groups (3.17 vs 3.0, p=0.78) (Figure 4i). This was graded using an accepted method used in previous hernia studies (33). In the SRT-PP group, one rat had grade 2 adhesions and one had grade 5 (colonic serosa was adherent to the mesh), whereas the rest had grade 3 (Figure 4b-d). All of the control rats had grade 3 adhesions (Figure 4f-h).

**Figure 4.**
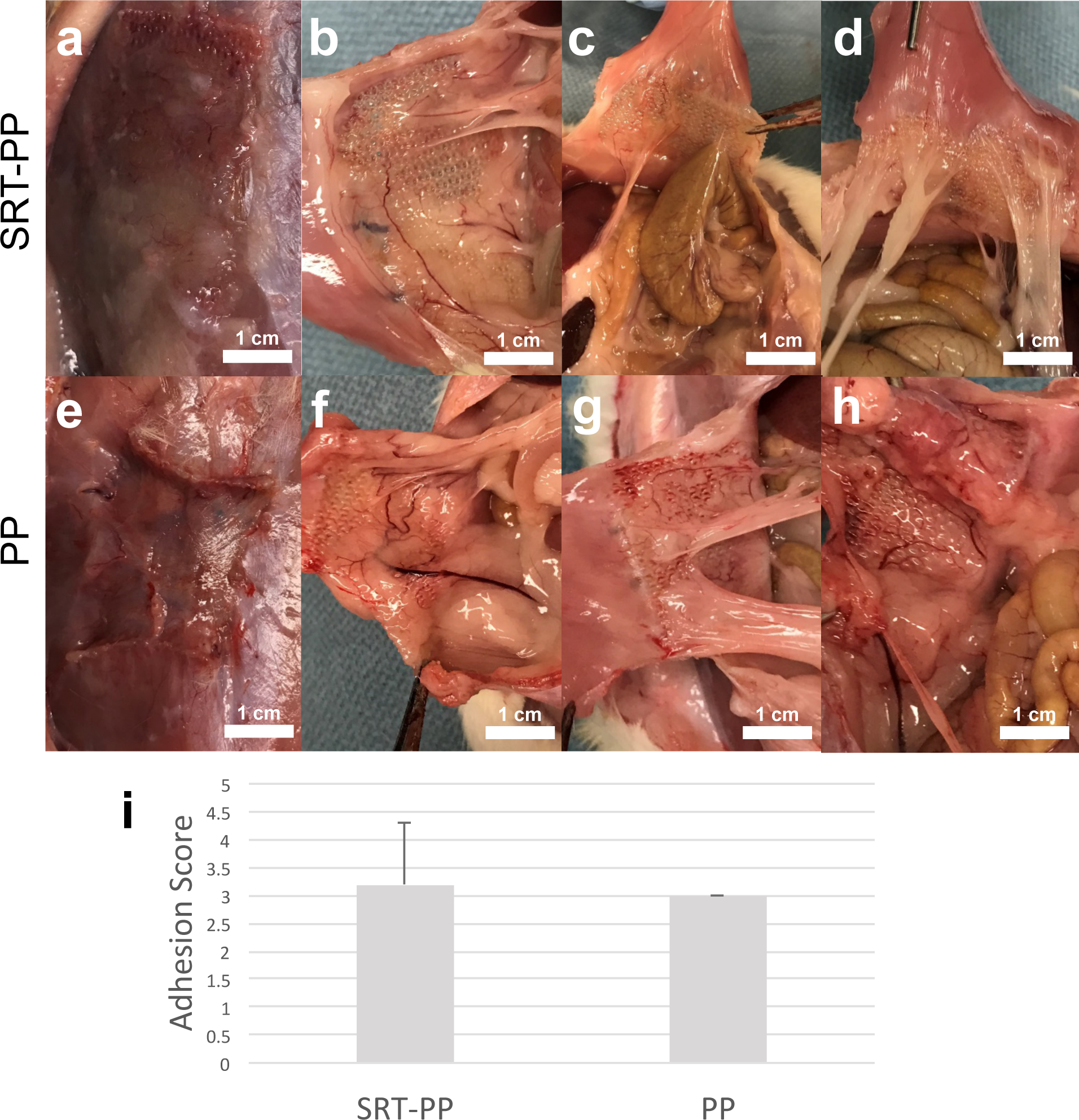
Necropsy and adhesion scoring. (a) SRT-PP rat at necropsy showing no contraction. (b) SRT-PP rat with grade 3 adhesions. (c) SRT-PP rat with grade 5 adhesion (colonic serosa attached to mesh). (d) SRT-PP rat with grade 2 adhesions. (e) PP rat at necropsy showing vertical mesh contraction. (f-h) PP rats with grade 3 adhesions. (i) Qualitative adhesion scores for composite SRT-PP and control PP.

### Histology

We performed H&E staining and microscopic assessment to show that ***c***hronic lymphoplasmacytic inflammation was identical in both groups (SRT-PP 3.3 vs PP 3, p=0.4) (Figure 6a,b,d,&e). Congruent inflammation between groups seemed to indicate a foreign body reaction to the mesh without any adverse effects from the SRT coating. Despite similar inflammation parameters, the SRT-PP group demonstrated increased fibrosis and collagen deposition ventral to mesh placement as demonstrated on trichrome staining (SRT-PP 168 µm vs PP 52 µm, p=0.005) (35). This appeared to confirm our *in vitro* study of cell proliferation where SRT coated meshes were characterized by a 3-fold increase in cell proliferation compared to uncoated PP mesh. We did not observe any systemic foreign body reaction and liver histology demonstrated normal architecture (Figure 5c&f) in both groups.

**Figure 5.**
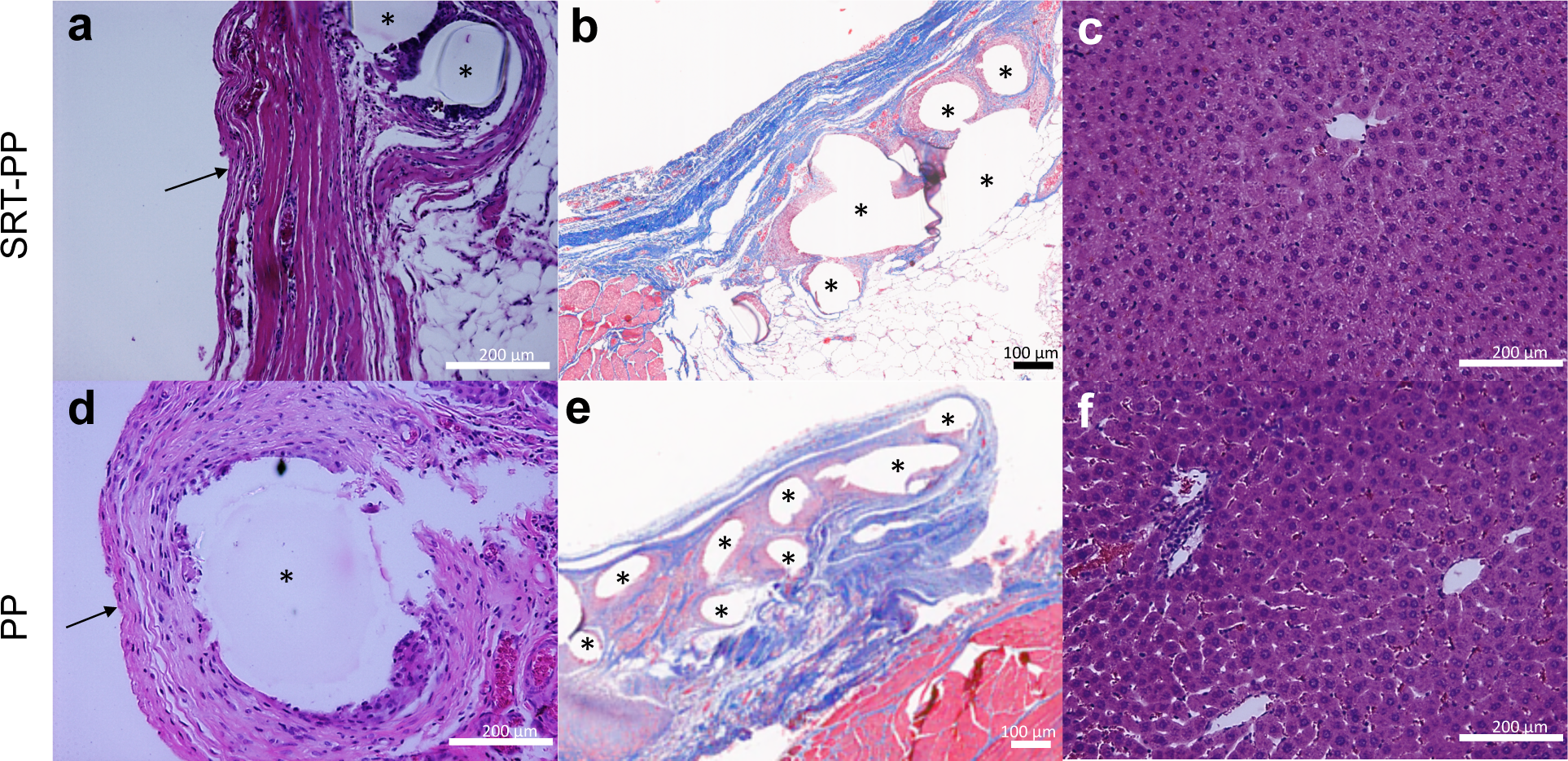
Histology. (a) H&E staining of SRT-PP rat abdominal wall+mesh (40X). (b) Trichrome staining of SRT-PP rat abdominal wall+mesh (10X). (c) H&E staining of SRT-PP rat liver (40X). (d) H&E staining of PP rat abdominal wall+mesh (40X). (e) Trichrome staining of PP rat abdominal wall+mesh (10X). (f) H&E staining of PP rat liver (40X). Arrows pointing to anterior fibrosis and asterisks (*) showing mesh fibers.

**Figure 6.**
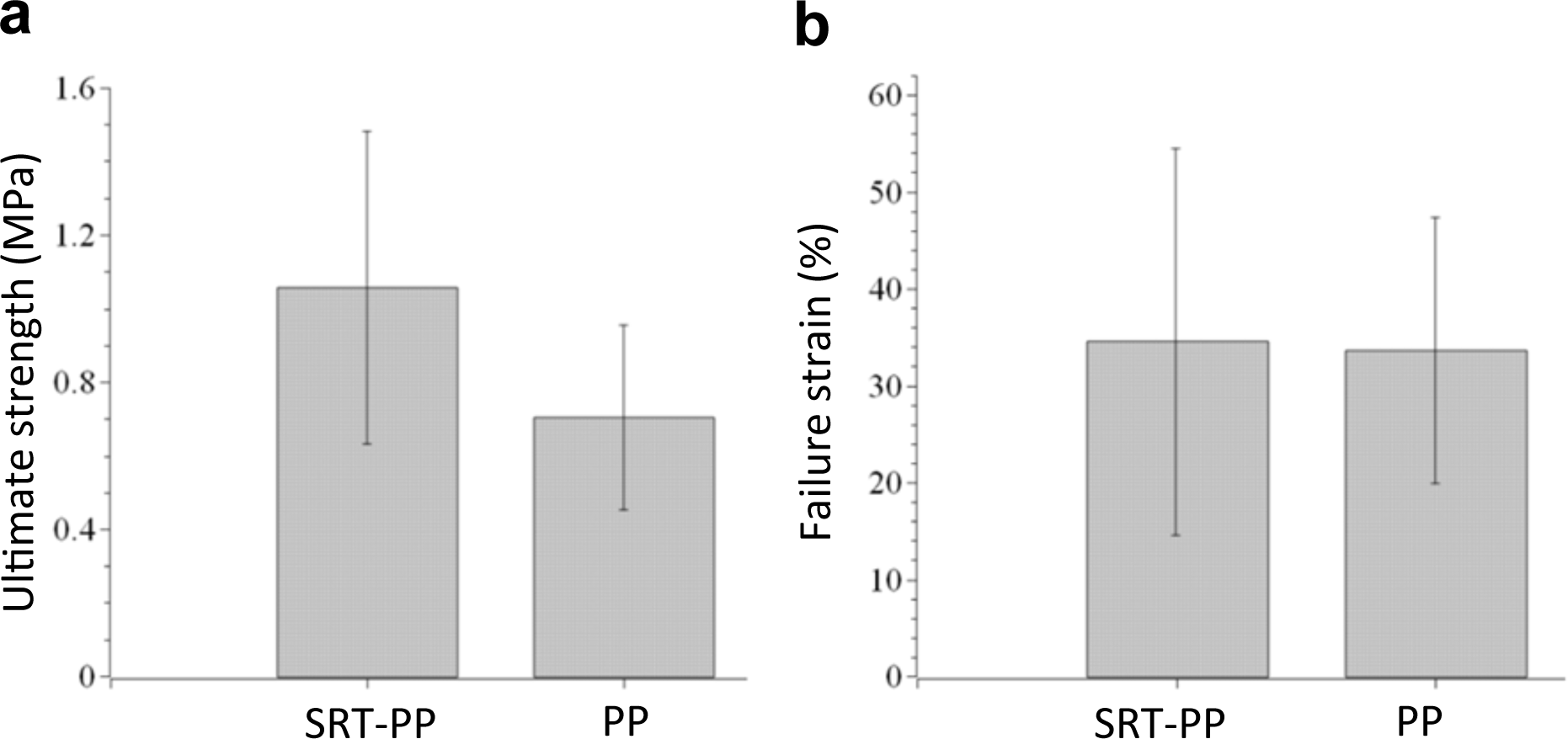
Biomechanical testing of explants. (a) Ultimate strength and (b) Failure strain of tissue explants seized from rats that have composite SRT-PP and control PP mesh.

### Biomechanical Testing of Explants

We performed biomechanical testing of mesh-tissue explants. The mechanical studies showed that tissue explants from SRT-PP mesh had greater adhesion strength (0.99 MPa vs 0.82 MPa, p=0.61) than the PP group. Additionally, SRT-PP tissue explants had greater failure strain (41% vs 33%, p=0.62). These are due to the increased integration at the mesh tissue interface in the SRT-PP group (i.e., SRT protein coating increases the integration). Although the small size of animal group tested in study, these results were statistically significant as shown in Figure 6.

## DISCUSSION

In our study, we demonstrated that a thin SRT coating increases the strength of the mesh-tissue interface in an inlay repair without any increase in inflammation. The SRT protein binds with abdominal wall proteins, while not relying on the typical inflammatory process of scar formation (36). High mechanical properties (i.e., elastic modulus of 2-4 GPa and strength of 50-100MPa) and enhanced tissue integration of SRT further helped to result in significantly less cranio-caudal shrinkage compared to uncoated PP mesh. Inflammation is known to increase tissue fibrosis across a wide spectrum of implants (37) fostering material embrittlement and fragmentation (38). These unique properties of SRT allow it to function better than other synthetic and biologic meshes, which can experience significant shrinkage (39) as demonstrated in both animal studies and clinically (15). Clinically, mesh shrinkage has been associated with significant post-operative pain in patients (40).

Furthermore, SRT demonstrated the ability to enhance abdominal wall integration when utilized as a composite coating on PP mesh. Histologically this appeared as an increase in scar formation of the anterior abdominal wall and was corroborated by increased ultimate strength. This collagen rich scar formation leads to the increase in mechanical strength as has been previously cited for other meshes (41). These characteristics allowed for similar complication rates, with regard to visceral adhesions, when compared to PP mesh.

These findings show that the SRT coating can be used to increase mesh tissue integration without any significant adverse effects, both locally and systemically. Although the material strength of synthetic mesh is adequate to restore abdominal wall integrity, either a sublay or onlay repair (Supplemental 4) is required to protect the viscera from the unwanted inflammatory effects. This leads to a paucity of durable options in patients where fascial re-approximation is not possible. For this patient population, SRT material may offer the most benefit. A biologic or Vicryl material coated with SRT or a pure SRT mesh could permit definitive fascial expansion with physiologic strength while not exacerbating concerns with visceral adhesions or erosion.

## CONCLUSION

This first time animal study showed that high-strength SRT protein is biocompatible. In addition, composite SRT-PP mesh showed superior abdominal wall integration compared to standard PP mesh. Since SRT is a biologically-derived material and provides high strength, it may prove useful in hernia repair. Its robust tissue integration without development of significant inflammation could permit for bridge implantation and definitive repair in patients where fascial re-approximation is not possible. When combined with its relatively low cost of production via industrial bio-fermentation compared to mesh extracted from animal tissues, it offers potential as either a mesh coating or a standalone mesh material. In either scenario, it is poised to significantly improve the biologic mesh armamentarium. Further testing can be conducted to assess the applicability of a pure SRT mesh and its local performance and systemic effects, which may change when a higher quantity is implanted. A rat model was chosen for our study as it limited research costs while allowing for the acquisition of preliminary animal implantation data for the first time. Future studies will evaluate SRT applicability in larger animals, such as swine, which provides better approximation to human abdominal wall strain and function.

## ACKNOWLEDGEMENTS

This work was partially supported by the Penn State College of Medicine (DR), and Penn State Materials Research Institute and Huck Institutes of Life Sciences (MCD and ITO). Research on fundamental properties of SRT protein is supported by Army Research Office under grant No. W911NF-16-1-0019 (MCD). The authors would like to thank the Turkish Ministry of National Education graduate scholarship (BA) and the International Postdoctoral Research Scholarship Program (BIDEP 2219) of the Scientific and Technological Research Council of Turkey (VO). The authors would also like to thank Aaron Selnick for helping in extraction of SRT rings, John Reibson (Penn State College of Medicine) for EtO sterilization of mesh, Donna Sosnoski (Penn State University) for her assistance in cell culture, and Dr. Jian Yang, PhD (Penn State University) for providing facilities for mechanical testing of explants. Additionally, Emma Dahmus, MSIII (Penn State College of Medicine) for her assistance in adhesion scoring, Jian-Li Wang, MD, PhD (Penn State College of Medicine) for his assistance in MRI interpretation, and Erik Washburn, MD (Penn State Health) for his assistance in histological assessment.

## CONFLICT OF INTEREST

The authors have issued and pending patent applications on SRT proteins.

